# Optical single-channel recording via diffusional confinement in membrane tethers

**DOI:** 10.1101/2025.05.07.652649

**Authors:** Madeleine R. Howell, Adam E. Cohen

## Abstract

Single-channel electrophysiology probes ion channel gating, but how can one probe membrane transport when the single-unit current is undetectable? We pulled membrane tethers from live cells to isolate individual transmembrane proteins. The tether constrained diffusion of transported substrate to the tether axis, leading to ∼1000-fold enhancement of substrate concentration and observation time compared to planar membranes. Fluorescent reporters inside the tether revealed individual transport events. We imaged unitary Ca^2+^ transport events in tethers containing the low-conductance T-type Ca^2+^ channel Ca_V_3.2, and compared our results to ensemble electrophysiology and stochastic gating simulations. This work establishes tether-based single-channel recordings as a powerful tool to study dynamics of membrane transport.

**TOC Graphic:** 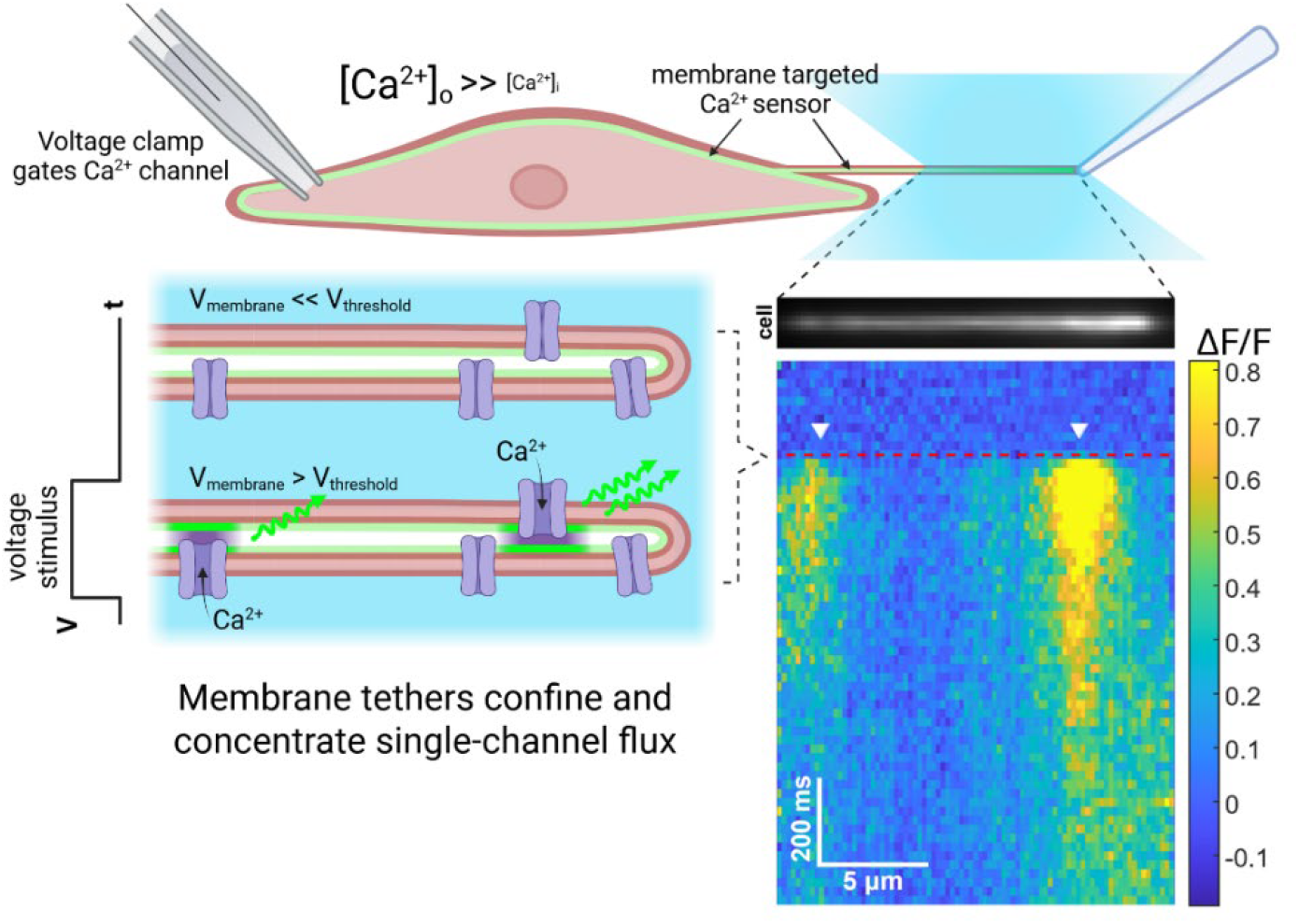

---

The plasma membrane of eukaryotic cells is decorated by diverse ion channels, transporters, and membrane-associated enzymes which orchestrate electrical and chemical signaling, osmotic regulation, and aspects of metabolism.^1–3^ The transport properties of these proteins are critical to their function. Single-channel electrophysiology^4–7^ can reveal molecular mechanisms of ion channel gating and regulation, but many channels and transporters remain inaccessible to this technique because either they are not electrogenic, or the single-unit current is too small (< ∼100 fA).^8,9^

Optical recordings of transmembrane flux using fluorescent reporters^10^ can also probe single-channel transport. Optical recordings have measured Ca^2+^ flux through single voltage- and ligand-gated channels, using either fluorogenic Ca^2+^ dyes or a genetically-encoded Ca^2+^ indicator (GECI) tethered to the channel.^11–18^ However, rapid dilution of transported ions into the cytoplasm limits the sensitivity of this approach. Isolation of single transporters in purified nanoscale proteoliposomes achieves single-channel sensitivity by preventing diffusive dilution of transported molecules.^19,20^ However, this cell-free approach requires protein purification, loses the native cellular context, and restricts molecular or electrical access to the lumen of the liposomes.

Membrane tethers consist of thin (typically ∼100 nm diameter) tubes of membrane which can be pulled from many cell types in culture.^21^ The tether lumen remains in diffusive interchange with the cell body, while the tether membrane restricts diffusion along the two orthogonal axes. Thus, tethers are intermediate between intact cell membrane and purified proteoliposomes: they provide substantial diffusive confinement but retain much of the physiological context and provide molecular and electrical access to the lumen.

Here, we pulled membrane tethers from intact cells and used a fluorescent reporter to record single-molecule gating events within the tether (Fig. 1A, B). We recorded single-channel events from Ca_V_3.2, a low-conductance (< 2 pS) and transiently activated voltage gated calcium channel (VGCC),^22,23^ using a membrane-targeted genetically encoded calcium indicator (GECI).^24,25^ Voltage-clamp in the parent cell controlled the gating of single channels. We related the single-channel gating propertes to ensemble Ca_V_3.2 currents, and used stochastic gating simulations to interpret our results.

**Figure 1.**
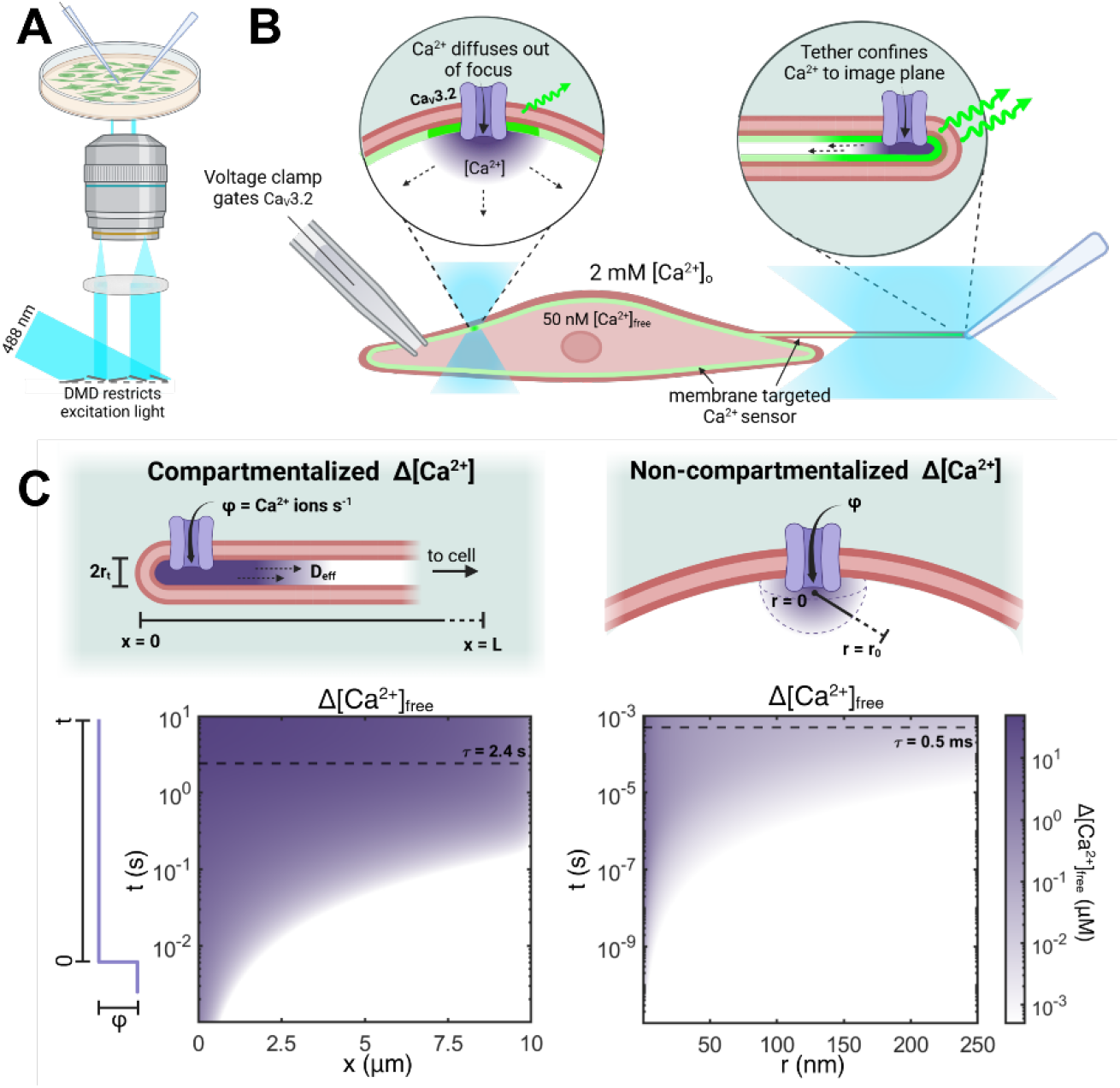
Single-channel recording in membrane tethers. **A**. Experimental setup comprising patterned illumination from a digital micromirror device (DMD) and two pipettes controlled by micromanipulators. **B**. Schematic of the experiment. A HEK293 cell expressed Ca_V_3.2 and a membrane-targeted Ca^2+^ indicator. One pipette applied voltage steps to gate the Ca_V_ channels. A second pipette pulled a membrane tether, which confined the Ca^2+^ influx from single Ca_V_ gating events. DMD-targeted illumination of the tether minimized background fluorescence and flare from the much brighter cell. **C**. Comparison of Ca_V_3.2 channel residing in a tether (left) or the cell body (right). Bottom: simulated Ca^2+^ profiles for the depicted geometries, assuming a step opening of the channel at *t* = 0, a transport rate *ϕ* = 10^5^ Ca^2+^ ions/s,^23^ an effective diffusion coefficient *D*_eff_ = 21 μm^2^/s,^32,46,47^ and a buffering capacity (buffer-bound Ca^2+^ per free Ca^2+^)^33^ κ = 200.^32,48^ In the tether, the channel is assumed to reside at the distal end. For the channel in the cell body, the Ca^2+^ concentration is averaged over a hemisphere of radius *r*. Dashed lines indicate the Ca^2+^ residence time in a tether-delimited (left) and diffraction-limited (right) focal volume. See *Supplementary Table 1* for diffusion model parameters.

## Results and Discussion

### Theory of single-channel signal enhancement in membrane tethers

We first modeled Ca^2+^ transport for a single channel residing either within a tether or on the cell body membrane. Details of the model are in *Supplementary Note 1* and *Supplementary Table 1*. We modeled the tether as a tube of radius *r*_*t*_ and length *L*, open to the cell body at its proximal end and sealed at its distal end (Fig. 1C). Tether radii were *r*_*t*_ ∼ 50 nm, set by the balance of membrane rigidity and membrane tension^26^ (Fig. S1), and lengths were *L ∼* 10 μm. We assumed rapid equilibrium of cytoplasmic Ca^2+^ buffers such that free Ca^2+^ flux, *ϕ*_free_, scales inversely with buffer capacity, *κ*_*B*_, following *ϕ*_free_ = *ϕ*/(1 + *κ*_*B*_), where *ϕ* is the total flux through the channel.^27^ The effective Ca^2+^ diffusion coefficient, *D*_eff_, is a weighted average of free and buffer-bound Ca^2+^ diffusion coefficients (Eq. 7 in *Supplementary Note 1*).^28^ We compared the Ca^2+^ concentration within a membrane-delimited tether volume to the mean Ca^2+^ concentration within a diffraction-limited confocal volume with radius *r*_*0*_, centered on a channel in the cell body membrane.

If a Ca^2+^ channel opens at the distal end of the tether, the steady-state free [Ca^2+^] concentration has maximal increase at the location of the channel:

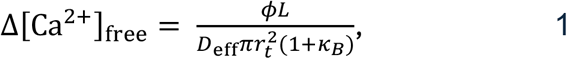

and a linear decrease down to Δ[*Ca*^2+^]_*free*_ = 0 at the junction with the cell body. The mean residence time of a Ca^2+^ ion in the tether is:

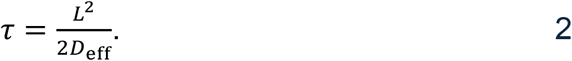

For a channel opening in the membrane on the cell body, we model the geometry as a planar membrane opening into an infinite half-space. This approximation is valid over distances much smaller than the size of the cell. In this scenario, Ca^2+^ rapidly diffuses into the open half-space (Fig. 1C). The steady state increase, Δ[Ca^2+^]_free_, decays with distance *r* from the channel as 1/*r*. We modeled the resulting Ca^2+^ domain as an average of Δ[Ca^2+^]_free_ over a diffraction-limited focal volume with radius *r*_*0*_ centered on the channel, so:

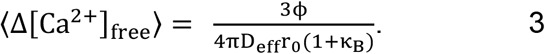

The mean residence time of a Ca^2+^ ion within this observation volume is:

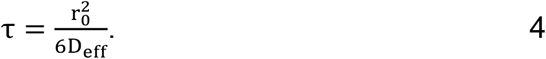

Our model predicts that a tether of typical dimensions (100 nm diameter, 10 μm long) enhances the in-focus Δ[Ca^2+^]_free_ near the channel by a factor of 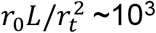 compared with diffraction-limited imaging of a Ca^2+^ channel in the membrane of the cell body (Fig. 1C, *Supplementary Note 1*). The time-resolution with which channel gating converts to changes in Δ[Ca^2+^]_free_ is ∼10^3^-fold slower in the tether than for the channel in the soma membrane. These simple estimates suggest that membrane tethers can render even miniscule transmembrane flows (*ϕ*_free_ = 100 s^-1^, *D*_eff_ = 200 μm^2^/s) detectable by a fluorescent reporter (K_D_ ∼ 1 μM).

### Tether fluorescence reports membrane Ca^2+^ influx with high fidelity

We tested whether Ca^2+^ compartmentalization in tethers permited optical recordings of single-channel gating. In a HEK293 cell line stably expressing doxycline-inducible Ca_V_3.2 α1 subunit^29^, we titrated doxycycline to achieve an expression density of ∼4 channels/μm^2^ (Fig. S2, Methods). We also expressed a membrane-targeted GECI, either lck-jGCaMP8f or GCaMP6s-CAAX^24,25^. We used a patch pipette in whole-cell mode to modulate the membrane voltage and to record ensemble Ca_V_ currents, and a second pipette to extract tethers. We used a digital micromirror device (DMD) to target 488 nm excitation light onto the tether and onto a small patch of the cell body opposite the junction with the tether (Fig. 2A). This illumination strategy minimized the total excitation light, decreasing background autofluorescence and flare from the bright fluorescence of the cell body, which might otherwise have overwhelmed the dim signal from the tether.

**Figure 2.**
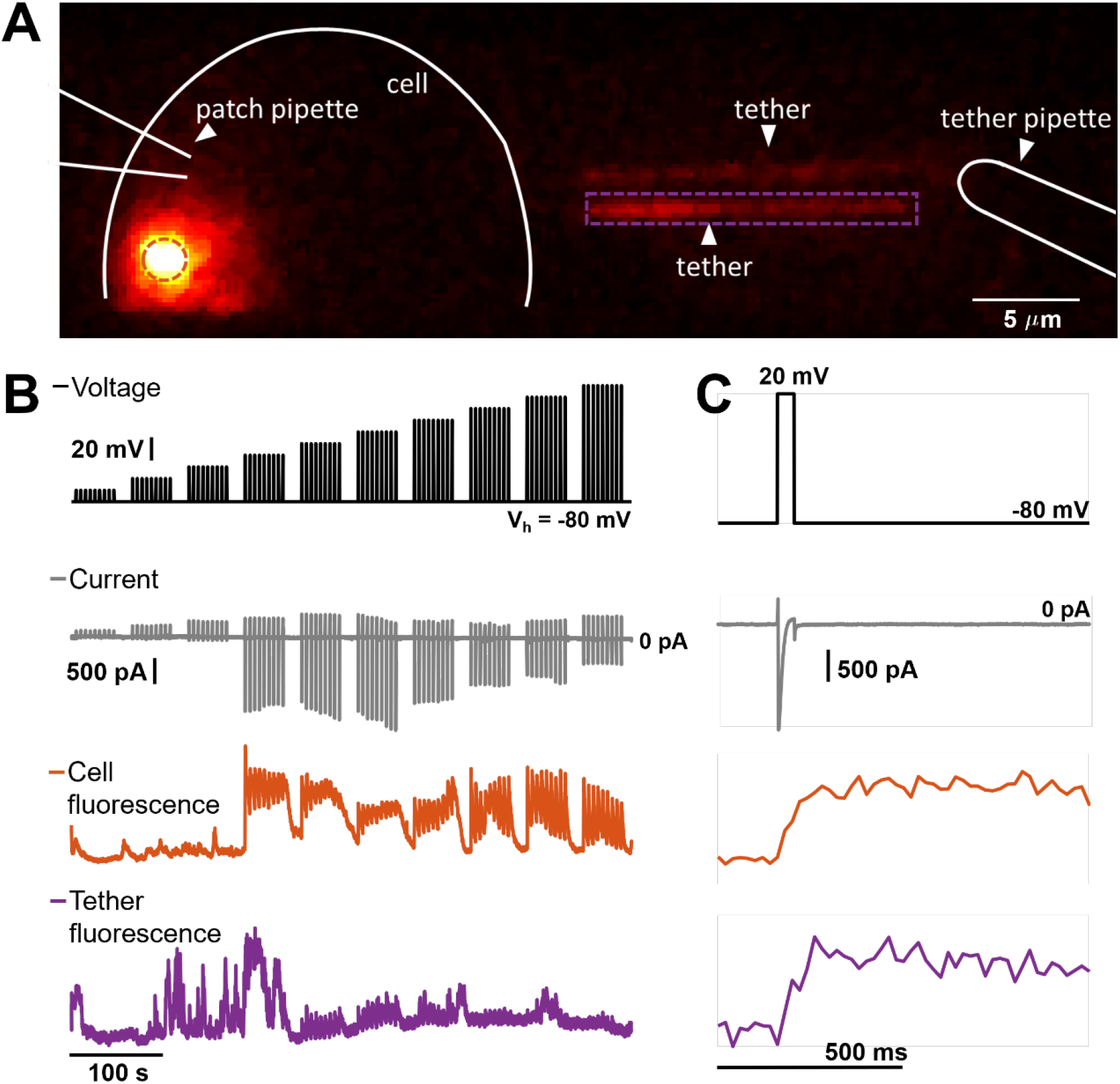
Simultaneous recording of tether and cell body Ca^2+^ dynamics. **A**. Ca_V_3.2 expressing HEK293 cell with localized excitation of a membrane-targeted GECI (GCaMP6s-CAAX) in the cell body (orange circle) and tether (purple rectangle). **B**. Concurrent patch-clamp recording of evoked currents (gray) and Ca^2+^-dependent fluorescence in the cell body (orange) and tether (purple) in response to depolarizing voltage steps (black) of 10-100 mV from a holding potential of -80 mV. **C**. Magnified view of traces in B. Trains of 9 repeats of 45 ms voltage pulses with a 5 s rest between pulses and a 15 s rest between pulse trains. Current and voltage traces were sampled at 100 kHz. Cell body and tether fluorescence were recorded at 50-200 Hz.

We applied trains of depolarizing voltage pulses from a holding potential of -80 mV, and simultaneously recorded whole-cell currents and GECI-reported Ca^2+^ dynamics in the cell, (ΔF/F)^C^, and tether, (ΔF/F)^t^ (Fig 2B-C). The camera frame-rate was 50-200 Hz. The voltage steps induced transient inward whole-cell currents which typically returned to zero before the end of the *t*_V_ = 45 ms voltage step (Fig. 3A). The time constants of Ca_V_3.2 activation and inactivation decreased for more depolarizing voltages, consistent with prior studies of this channel^22^.

**Figure 3.**
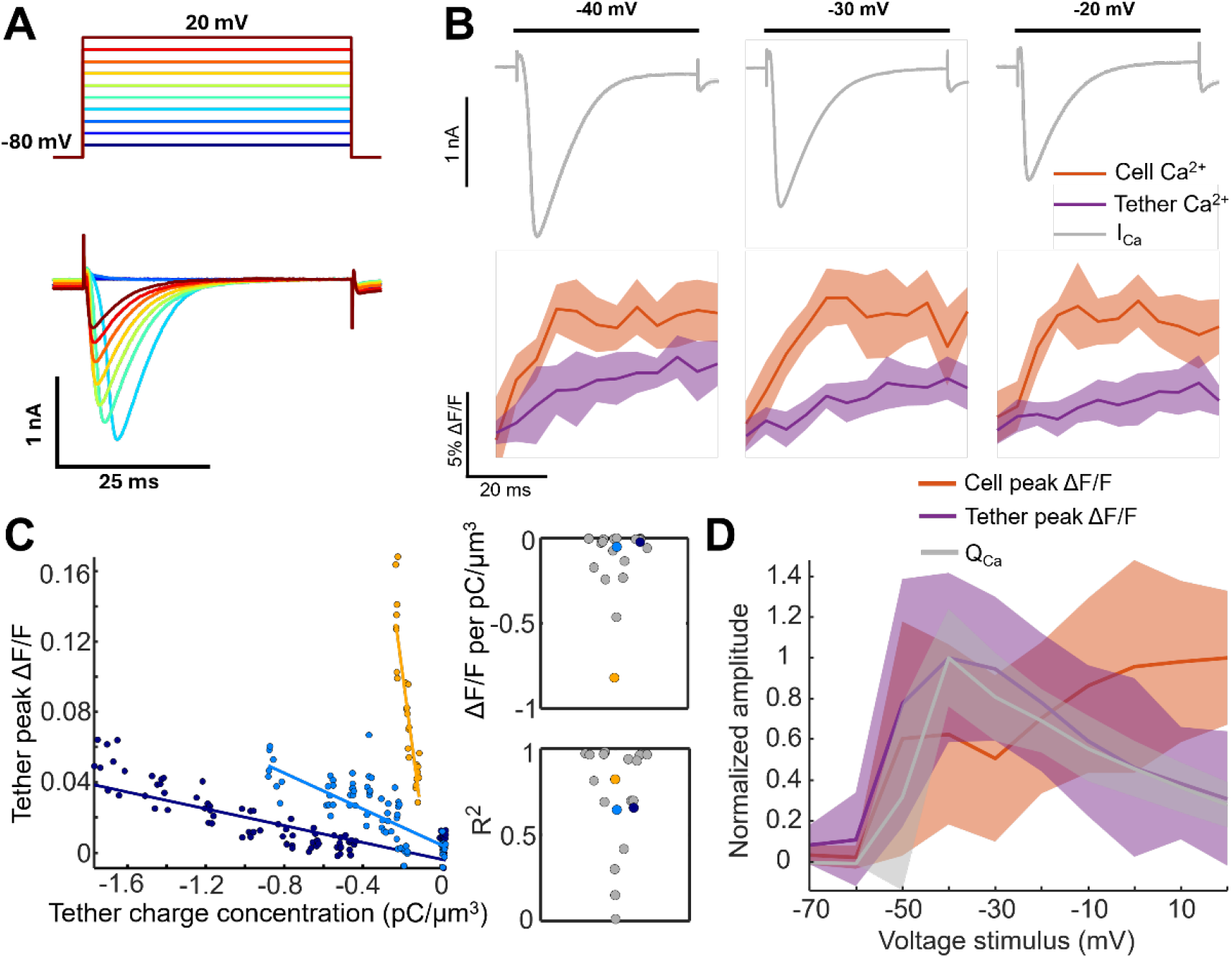
Tether Ca^2+^ signal correlates with whole-cell charge transport. **A**. Inward currents (bottom) recorded from a Ca_V_3.2 expressing HEK293 cell in response to 45 ms depolarizing voltage steps (top). **B**. Trial averaged current (top), cell (orange) and tether (purple) lck-jGCaMP8f fluorescence responses to membrane depolarizations. To calculate (ΔF/F)^t^, ΔF/F was first calculated for each position along the tether using the pre-stimulus baseline fluorescence as F, then averaged along the tether length. Data shown for one cell (n = 9 trials, std. dev. shading). **C**. Peak (ΔF/F)^t^ amplitude of tether lck-jGCaMP8f fluorescence transients as a function of predicted tether charge concentration, 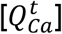, and linear fits. Data and fits shown for three example tethers. Inset: slope (top) and R^2^ (bottom) from linear fits of n = 20 tethers. **D**. Voltage dependence of stimulus-evoked charge influx (gray), tether (purple) and cell (orange) peak ΔF/F amplitudes averaged across n = 25 lck-jGCaMP8f cell-tether pairs. Traces are normalized for each tether-cell pair before averaging and shaded to show std. dev. Tether peak (ΔF/F)^t^ and Q_Ca_ were more closely correlated than were cell peak (ΔF/F)^C^ and Q_Ca_.

Membrane depolarization evoked Ca^2+^ signals in the cell and tether (Fig. 2B-C). Both regions also showed occasional spontaneous Ca^2+^ dynamics. An estimate of tether electrotonic length-constant, *λ*, based on whole-cell membrane conductance and tether geometry, gave *λ* ∼ 110 μm (membrane resistivity *R*_m_ = 1.5 × 10^6^ MΩ μm^2^; intracellular resistivity *R*_*i*_ = 3.0 MΩ μm;^30^ radius *r* = 50 nm), indicating that Ca_V_ channels throughout the tether experienced membrane voltages similar to the voltage at the cell body. The appearance of voltage-gated Ca^2+^ influx in the tether was consistent with this finding.

For both cell body and tether, the stimulus-evoked GECI fluorescence grew and decayed slowly compared to the patch-clamp recorded currents (Fig 2C, Fig. 3B). The upstroke of the GECI fluorescence was much faster (∼10 ms) than the recovery (∼1.4 s), implying that the peak GECI fluorescence was proportional to the increase in Ca^2+^ concentration due to the voltage step:

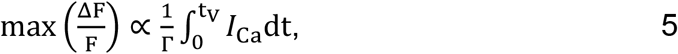

where *Γ* is the volume into which the Ca^2+^ is diluted and *I*_Ca_ is the absolute value of the inward calcium current.

For each voltage step, we calculated whole-cell charge influx, 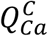, via the integral of the voltage-clamp current recording. We estimated the surface areas of the tether and of the whole cell from the tether geometry and the whole-cell capacitance, respectively. We assumed that channel density was approximately homogeneous across the cell and the tether, so the Ca^2+^ flux was apportioned between the cell 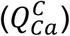 and the tether 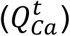 according to their relative surface areas.

We then checked the proportionality across voltage steps between tether Ca^2+^ signal, (ΔF/F)^t^, and the predicted tether charge concentration, 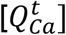. We observed a clear linear relationship within each tether (Fig. 3C, S3). The differing slopes between tethers likely reflected variation in channel density between tethers. We also observed trial-to-trial fluctuations about the linear relationship, which we attribute to fluctuations in channel occupancy and activation within tethers (Fig. 3C inset). These fluctuations are not unexpected, considering the small number (∼12) of stochastically gating channels per tether (Fig. S2).

Unexpectedly, we found that the correlation between (ΔF/F)^t^ and 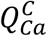 was stronger than the correlation between (ΔF/F)^C^ and 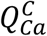 (Fig. 3D, S3), i.e. whole-cell charge and fluorescence-reported Ca^2+^ influx were not well correlated. We speculate that the Ca^2+^ in the cell body may have had additional stochastic uptake and release from internal Ca^2+^ stores (e.g. in mitochondria and endoplasmic reticulum), whereas these organelles were excluded from the tether. Since we could not establish a robust relationship between Ca^2+^ influx, Ca^2+^ concentration, and GECI fluorescence in the cell body, we were unable to establish an absolute relation between (ΔF/F)^t^ and 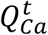 . Hence, subsequent measurements focused on analyzing relative changes in (ΔF/F)^t^.

### Ca^2+^ indicator reports discrete influx events

We observed stimulus-triggered Ca^2+^ events which originated from distinct positions along the tether and which fluctuated in number and position between successive voltage steps (Fig. 4A, S4). We visualized these Ca^2+^ events via kymographs, where fluorescence was displayed as a function of position along the tether and time. We combined smoothing and automated thresholding to identify peaks that were > 3 ± 0.9 *σ* from baseline fluctuations (Methods). By comparing recordings taken at the holding potential of -80 mV (when Ca_V_3.2 channels are expected to be closed) to the recordings during voltage steps, we estimate a false-positive rate of 0.9% (Fig. S5). We selected well-separated (> 7 μm) Ca^2+^ events and then calculated stimulus-triggered average spatio-temporal footprints (Fig. 4B, S4). These footprints captured the rapid depolarization-triggered Ca^2+^ influx and spread. Since the response-time of the Ca^2+^ measurements (∼40 ms to peak) was comparable to the 45 ms duration of the voltage pulse, we could not resolve variations in the onset of Ca^2+^ influx relative to the voltage upstroke. We observed fast and slow components to the fluorescence decay (Fig. 4C, S4), which we provisionally ascribe to local Ca^2+^ buffer equilibration and diffusional spreading, respectively.

**Figure 4.**
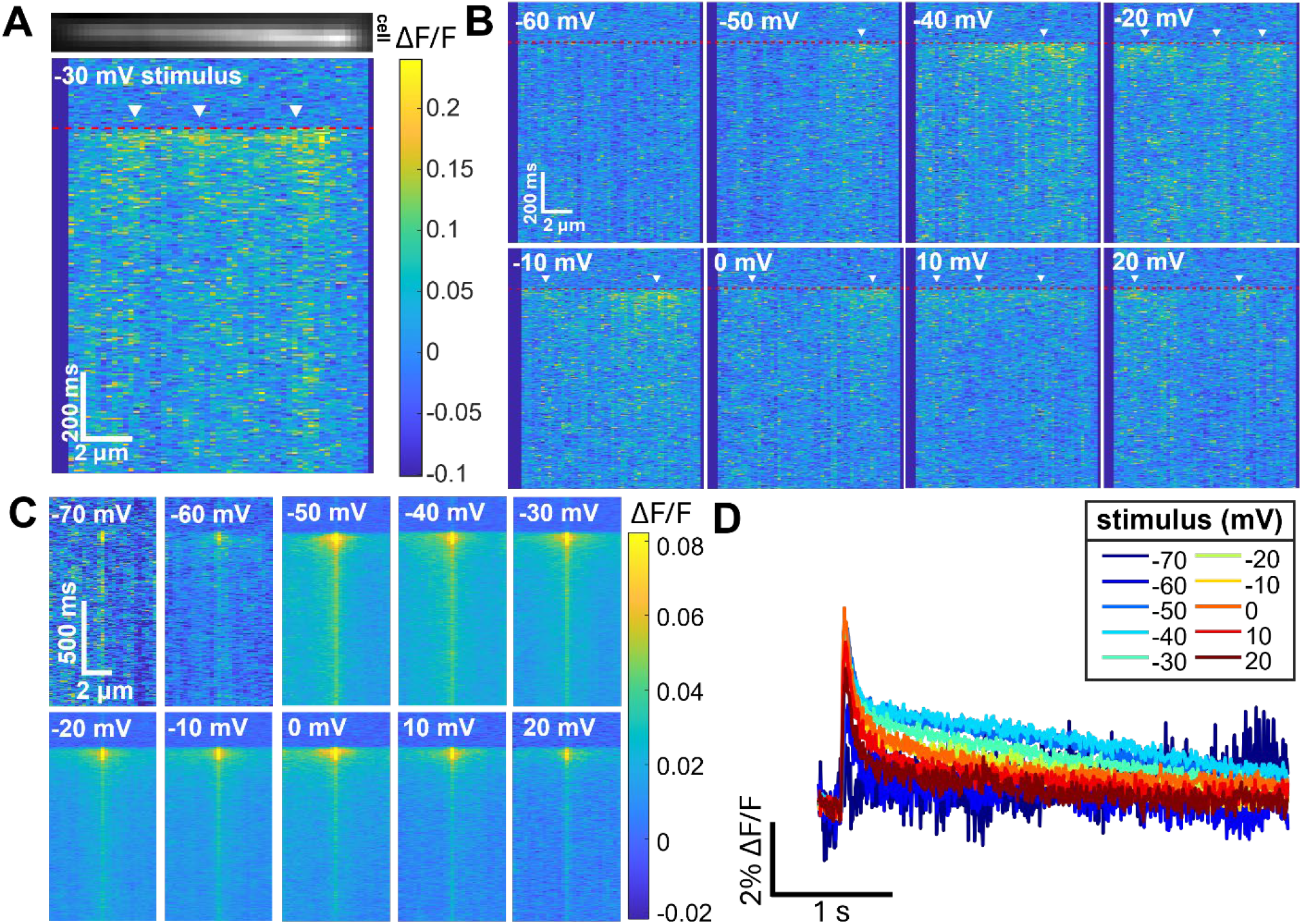
Tether Ca^2+^ signal comprises discrete events. **A**. Epifluorescence image of a tether above a kymograph for a 45 ms voltage step from -80 mV to -30 mV. Red dashed line indicates step onset. White arrowheads indicate discrete Ca^2+^ events. **B**. Kymographs of tether Ca^2+^-dependent fluorescence in response to voltage steps from - 80 mV to between -60 mV and +20 mV. Data shown for single-trial responses of one tether. **C**. Stimulus-triggered average kymographs of spatially isolated tether Ca^2+^ events (8-251 events per voltage, n = 25 tethers, 17 cells). **D**. Time-course of events in (C), averaged over space.

### Amplitude and frequency of single-channel gating events

We next compared event characteristics reported by tether lck-GCaMP8f to whole-cell electrical measures of channel gating. For each voltage step, we constructed a histogram of the unitary event amplitudes (ΔF/F)^t^ (Fig. 5A). We reasoned that the ensemble average of these discrete gating events should reproduce the previously observed linear relationship between (ΔF/F)^t^ and 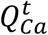 (Eq. 5). To test this idea, we summed all unitary event amplitudes for each voltage. The plot of cumulative (ΔF/F)^t^ vs. voltage closely followed the plot of 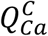 vs. voltage, determined from the whole-cell patch clamp measurements (Fig. 5B). Thus, the fluorescence-detected Ca^2+^ events, when combined to create an ensemble average, recapitulated the voltage-dependent behavior of the macroscopic currents.

**Figure 5.**
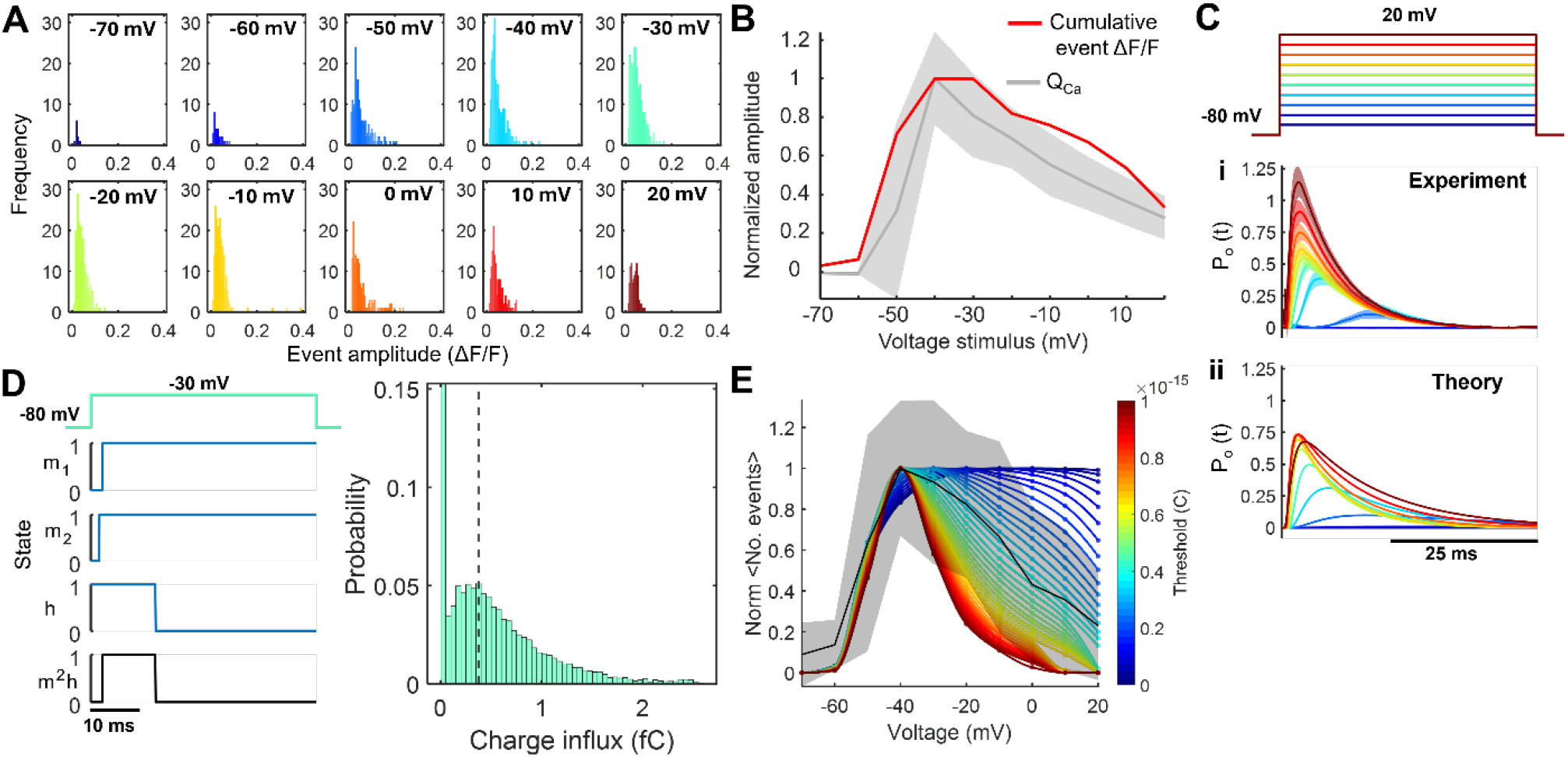
Single-molecule gating properties of Ca_V_3.2. **A**. Histograms of tether Ca^2+^ event amplitudes (10-277 events recorded per voltage stimulus, n = 25 tethers, 17 cells). **B**. Cumulative amplitudes of the event histograms in A (red) and average charge influx for the corresponding cells (gray) as a function of voltage (n = 25 cell-tether pairs). Shading indicates std. dev. **C**. Ca_V_3.2 channel open probability, P_O_, in response to depolarizing voltage steps from a holding potential of -80 mV (top). (i) P_O_ estimated from inward currents (n = 17 cells, shading indicates s.e.m.) and (ii) P_O_ calculated using a Hodgkin-Huxley-like model of gating transitions. **D**. Left: stochastic simulation of a channel gating trajectory in response to depolarization of duration *t*_V_. Blue traces show individual gate trajectories, black trace shows channel state (m^2^h). Right: distribution of charge influx, *q*, due to single-channel openings in response to depolarization to -30 mV for duration *t*_V_ (n = 10^4^ channel trajectories). Calculated by integrating Eq. 12 over *t*_*V*_. Black dashed line indicates predicted detection threshold from E. **E**. Frequency of observed events vs. voltage (normalized, black, shading std. dev. of n = 19 tethers) and predicted single-channel event probability for different detection thresholds (colors). Color bar indicates simulated charge detection threshold (maximum R-value at 0.38 fC threshold).

We then sought to model the distribution of (ΔF/F)^t^ at each voltage. This distribution is not directly accessible from ensemble-average patch clamp measurements. We thus turned to stochastic simulations of channel gating. We fit a Hodgkin-Huxley-like model of channel gating to the whole-cell currents recorded across voltage steps from -70 mV up to +20 mV (Fig. 5C). We then performed stochastic simulations of single-channel gating, using the kinetic parameters derived from the fit to the whole-cell currents (Fig. 5D, *Supplementary Table 2, Supplementary Notes 2*,*3*, Fig. S6). For each voltage step, we calculated the distribution of single-channel 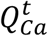 values, taking into account the voltage-dependent gating dynamics and the driving force for Ca^2+^ entry.

In the simulated single-channel trajectories, the probability of channel opening saturated at depolarized potentials (Fig. S6). In contrast, we observed a decrease in the frequency of Ca^2+^ events at depolarized potentials (Fig. 5E, black trace). At depolarized potentials, we expect faster Ca_V_3.2 channel kinetics, lower driving force for Ca^2+^ entry and, thus, lower unitary charge passed by the channel. We hypothesized that these very small Ca^2+^ influx events might be below our detection threshold. To test this hypothesis, we applied a threshold to the simulated distributions of 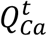 (Fig S6) and counted the voltage-dependent frequency of events that exceeded this threshold. Upon varying the simulated detection threshold, we found best correspondence between simulation and experiment at a threshold of 0.38 fC for GCaMP8f and 0.4 fC for GCaMP6s (Fig. 5D,E, S7). This detection threshold corresponds to an equivalent constant current of ∼9 fA over *t*_*V*_ = 45 ms; or to 100 fA for a 4 ms gating event. These unitary charges are substantially smaller than are typically recorded by patch clamp.^6,7^

## Conclusion

Historically, it has been challenging to measure small ionic or biomolecular fluxes in intact membranes. Here, we showed that membrane tethers can compartmentalize and thereby amplify single-channel fluxes. Using membrane-targeted GECIs in tethers, we detected puncta of elevated Ca^2+^ in response to voltage-dependent gating of a low-conductance Ca_V_3.2 channel (Fig. 4). Despite slow GECI kinetics, the tethers amplified and prolonged [Ca^2+^] transients enough that single-channel gating events were readily detectable (Fig. 1).

We compared the amplitude and frequency of tether Ca^2+^ events to ensemble Ca_V_3.2 electrophysiology. The mean height of tether Ca^2+^ transients correlated with the charge passed during depolarizing stimuli (Figs. 3, 5). Conversely, cell GECI fluorescence was poorly correlated with transported charge, likely due to intracellular Ca^2+^ transport. By comparing simulated single-channel gating trajectories and the observed frequency of Ca^2+^ events, we estimated a detection sensitivity of ∼0.4 fC, corresponding to ∼1250 Ca^2+^ ions (Fig. 5). Strong buffering of intracellular Ca^2+^ presented a challenge for our measurements. It is estimated that the basal buffering capacity of cytoplasm ranges from 100 to 200,^31–33^ so an influx of 1250 Ca^2+^ ions corresponds to only 6 – 13 free Ca^2+^ ions.

In principle, our tether-based recording scheme can generalize to other proteins (e.g. channels, transporters, membrane associated enzymes). The key requirement is a suitable fluorescent reporter of the substrate or products^10^. For a 10 μm tether, a turnover rate of 100 s^-1^ at the distal end of the tether, and an unbuffered substrate with a diffusion coefficient of 200 μm^2^/s, the local concentration increase is 1 μM, and the substrate residence time is 250 ms. These parameters benchmark the sensitivity needed for different substrate fluxes. Many parameters contribute to determining flux through a transport protein, including the chemical and electrical potentials of transported species, permeation mechanism, protein conformational dynamics, bilayer tension, post-translational modifications, cofactor binding, and redox chemistry.^1–3^ Single-channel recordings in tethers open the door to exploring these factors in a cellular context.

Some cellular membranes natively contain nanoscale compartments, such as dendritic spines^34^, neurites^35,36^, primary cilia^37^, synaptic vesicles^38^, tunneling nanotubes^39^, and retraction fibers.^40^ Our work highlights the possibility that stochastic single-channel gating events within these structures can lead to substantial fluctuations in substrate concentration. Whereas membrane signaling is typically modeled via ensemble-average kinetics, a direction for future research will be to explore the biological roles of stochastic single-molecule gating in nanoscale compartments.

## Methods

### Genetic constructs

We used membrane targeted Lck-jGCaMP8f and GCaMP6s-CAAX Ca^2+^ indicators to detect single-channel tether Ca^2+^ transients. Both constructs were expressed under the control of a CMV promoter. The *pGP-CMV-GCaMP6s-CAAX* construct was a gift from Tobias Meyer (Addgene #52228) and used as provided. The *pGP-CMV-lck-jGCaMP8f* construct used in this work was made from a *pZac2*.*1-GfaABC1D-lck-jGCaMP8f* plasmid gifted by Loren Looger (Addgene #176759).

*lck-jGCaMP8f* was cloned into a pGP-CMV vector backbone (Addgene #104483) using Gibson assembly. Briefly, the vector was linearized by sequential digestion using restriction enzymes (New England Biolabs) and purified by GeneJET gel extraction kit (ThermoFisher). The insert fragment was generated by polymerase chain reaction amplification and inserted into the backbone using NEBuilder HiFi DNA assembly kit (New England Biolabs). The resulting construct was verified by sequencing (Primordium).

### Cell culture and transfection

HEK293 cells stably expressing Tet repressor (T-REx-293, ThermoFisher), constitutive human K_ir_2.3, and doxycycline-inducible human Ca_V_3.2 were a generous gift from Terrance Snutch. Ca_V_3.2/K_ir_2.3 cells were maintained at 37 °C and 5% CO_2_ in Dulbecco’s modified Eagle’s medium (DMEM) formulated with high glucose, GlutaMAX, and pyruvate (Cat. No. 10569010, ThermoFisher) and supplemented with 10% heat-inactivated fetal bovine serum, penicillin (100 U/mL), and streptomycin (100 µg/ mL). Culture medium was supplemented with geneticin (600 μg/mL, Life Technologies), hygromycin B (150 μg/mL, Sigma Aldrich), and blasticidine (10 μg/mL, ThermoFisher) selection agents. Selection agents were removed and cells were passaged at least 24 hr prior to transfection with GECI constructs. Passaged cells were grown to ∼80% confluence prior to transfection with TransIT-293 (Mirus Bio) according to manufacturer protocols. Ca_V_3.2 expression was induced 24-72 hr prior to imaging using 1.5 ng/mL doxycycline (Sigma). This doxycycline dose generated a whole cell Ca_V_3.2 conductance of 5.6 ± 1.5 pS/μm^2^ (n = 17 cells) at -20 mV (Fig. S2E).

### Patch-clamp electrophysiology

Whole cell voltage-clamp recordings were acquired from Ca_v_3.2 expressing cells. On the morning of an experiment, cells were trypsinized and replated on poly-D-lysine (Sigma) coated glass bottom dishes (20 μg/mL incubated at RT overnight or 20 min at 37 °C, washed 3x with phosphate-buffered saline). Prior to an experiment, cells were washed 2x and immersed in an extracellular solution containing (in mM): 125 NaCl, 25 glucose, 15 HEPES, 2.5 KCl, 1 MgCl_2_, 2 CaCl_2_. Patch pipettes (1-10 MΩ) were filled with an internal solution containing (in mM): 8 NaCl, 130 KMeSO_3_, 10 HEPES, 4 MgATP, 0.3 Na_3_GTP, 5 KCl. The pH was adjusted to ∼7.3 using KOH and the osmolarity was adjusted to ∼295 mOsm/L with sucrose. Signals were amplified using an Axopatch 200B amplifier (Molecular Devices), filtered at 5 kHz and digitized at 100 kHz (DAQ PCIe-6323, National Instruments). A micromanipulator (Sutter) maneuvered patch pipettes to the cell membrane. Cell membrane voltage was clamped at a holding potential of -80 mV and then stimulated with depolarizing pulse trains (9x 45 ms pulses, 5 s inter-pulse rest, 15 s inter-train rest, Fig. 2B-C). All experiments were performed at 32 °C with temperature control provided by an objective heater (Bioptechs) and stage heater (Warner).

### Tether formation

Micropipettes were pulled from glass capillaries (World Precision Instrument, 1B150F-4) using a pipette puller (Sutter P1000) and the tip of the pipette was sealed and rounded using a microforge (WPI, DMF1000) to form a microneedle. Microneedles were surface treated via incubation in poly-D-lysine (100 μg/mL, 37 °C for 20 min) followed by incubation in concanavalin A (100 μg/mL, Vector Laboratories) until use. A micromanipulator maneuvered the microneedle to the cell surface and tethers were formed upon brief contact and retraction of the microneedle tip from the membrane.

### Imaging

Ca^2+^ imaging was performed on a custom-built inverted microscope using a 60x water immersion objective (Olympus, UPLSAPO60XW). Illumination, spatial light patterning, patch amplifier, data acquisition card, and camera were controlled with custom MATLAB (Mathworks) based software^41^. A digital micromirror device (DMD, Texas Instruments DLP3000) restricted 488 nm excitation light (81 mW/mm^2^, PhoXX-488-60) to cell and tether ROIs. GECI fluorescence was imaged onto an EMCCD camera (DU-897E-CSO-#BV, Andor) with 300x electron-multiplying gain at a rate of 50-200 Hz. An adjustable slit (VA100, Thorlabs) restricted incident light to a subset of camera rows to enable high-speed imaging. Camera frames were either acquired at the rising edge of patch-clamp voltage stimuli or aligned in post-processing. Fluorescence recordings were registered in time to the voltage-clamp data by reference to the shared the DAQ clock.

### Image processing and Ca^2+^ event finding

Tether orientation and transverse offset were identified using the Random Sample Consensus (RANSAC) algorithm.^42^ Tether motion was compensated in post-processing to stabilize the image of the tether. Fluorescence was averaged along the width (short axis) of the tether and plotted as a function of time and position along the length of the tether. To these kymographs, we applied a median filter and a spatio-temporal smoothing filter. We calculated a global threshold for each recording using Otsu’s method^43^. To segment Ca^2+^ events, we thresholded each recording and assigned peak amplitude and position to local maxima^44^. In instances where events were detected less than 3.5 μm apart, the smaller amplitude event was discarded. Additionally, we discarded any kymograph frames with greater than 0.5 events/μm.

### Data Analysis

Data processing and image analysis were performed in MATLAB (Mathworks) and ImageJ^45^. Ca^2+^ diffusion modeling is described in *Supplementary Note 1*. Ca_V_3.2 channel modeling and stochastic simulations are described in *Supplementary Notes 2*,*3*. To compare observed Ca^2+^ event frequency to that predicted by our stochastic simulations (Fig 5D,E), we first calculated the mean events vs. voltage per tether and then averaged across tethers. Then, starting with the simulated distributions of charge influx, we applied a charge detection threshold, and calculated the expected frequency of detected events vs. voltage. We varied the simulated detection threshold to identify the threshold that gave closest correspondence to our data.

## Supporting Information

Contains ensemble electrophysiology data for Ca_V_3.2 expressed in HEK293 cells, whole-cell and single-channel fluorescence recordings of Ca^2+^ influx using GCaMP6s-CAAX, and descriptions of Ca^2+^ diffusion model, Ca_V_3.2 gating model, and stochastic simulations.

## Acknowledgements

We thank A. Preecha, S. Begum, and C. Bodden for technical assistance, S. Innes-Gold, H. Davis, and F.P. Brooks for assistance with instrumentation, P. Park and Y. Wang for assistance with electrophysiology and E. Moult and K. Xiang for helpful discussions regarding data analysis and modeling. We thank T. Meyer, L. Looger, and D. Kim for GECI plasmid constructs and T. Snutch for inducible Ca_V_3.2 cell line. This work was supported by Vannevar Bush Faculty Fellowship N00014-18-1-2859 (A.E.C.). and National Science Foundation Graduate Research Fellowship Grant #DGE 2140743 (M.R.H.). Figs. 1, S1C, S2A and TOC graphic contain elements generated in BioRender.

---

**Supplementary Note 1. Model of single-channel Ca**^**2+**^ **domains in a tether and cell body membrane**. We assume: 1) The equilibration time of Ca^2+^ buffering is fast relative to diffusion out of the observation volume, so free and buffer-bound Ca^2+^ exist in a local equilibrium. 2) The change in total (free + bound) Ca^2+^ concentration, Δ[Ca^2+^]_T_, due to a single channel gating event does not saturate endogenous buffers. Therefore, the change in free Ca^2+^, Δ[Ca^2+^]_free_, can be expressed as a fraction of Δ[Ca^2+^]_T_:

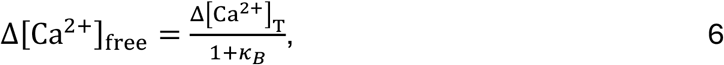

where *κ*_*B*_ is the cytoplasm Ca^2+^ buffering capacity.

The effective diffusion coefficient of Ca^2+^ is a weighted average of free (D_Ca_) and buffer-bound (D_B_) Ca^2+^ diffusion coefficients:

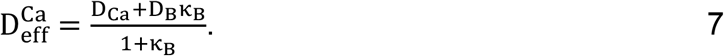

To calculate the spatiotemporal profiles of single-channel Ca^2+^ domains in a tether, we solve the diffusion equation numerically using the MATLAB *pdepe* solver,

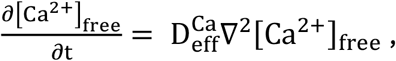

with boundary conditions:

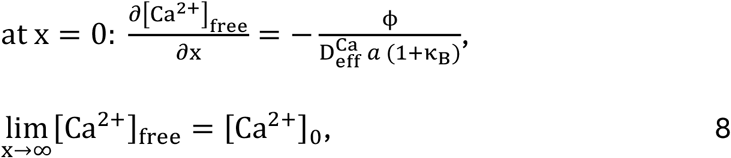

where *x* denotes distance from the open channel and *a* denotes tether cross-sectional area. Δ[Ca^2+^]_free_ profiles are calculated using the model parameters in Table S1. We used the same diffusion model to calculate volume-averaged Δ[Ca^2+^]_free_ in the cell body.

**Supplementary Note 2. Model of T-type voltage-gated Ca**^**2+**^ **channels**. We employ the Hodgkin-Huxley (HH) model of T-type Ca_V_ currents developed by Huguenard and McCormick^*49*^. Briefly, the macroscopic current, *I*_*Ca*_ is defined as the product of the unitary Ca^2+^ conductance, *g*_*SC*_, the ensemble-averaged channel open probability, *P*_*O*_, the number of Ca_V_ channels in the membrane, *N*_*C*_, and the voltage and [Ca^2+^]-dependent driving force, *G*.

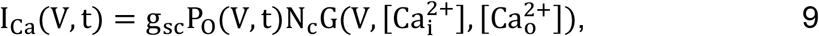

where *P*_*O*_ is modeled as the product of three independent two-state gates: two activation *m*-gates and one inactivation *h*-gate.

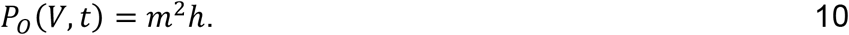

To fit *P*_*O*_ parameters (see below, ***Thermodynamic model of rate constants***) to our ensemble currents, we used *G* defined by the Goldman-Hodgkin-Katz current equation (in units of C m^-3^) and a membrane permeability (*P*, m^3^ s^-1^), such that

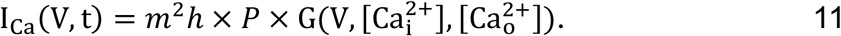

For stochastic simulation of single-channel trajectories (see *Supplementary Note 3*), we define the unitary current, *i*_*Ca*_, as the product of *P*_*O*_, *G*, and a single-channel permeability, *p* (same units as above).

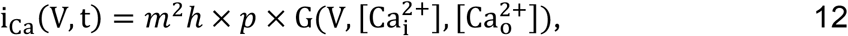

where *p* is related to the single-channel slope conductance^*50*^ *g*_*sc*_, measured by *Weber et al*. in the linear i-V regime:

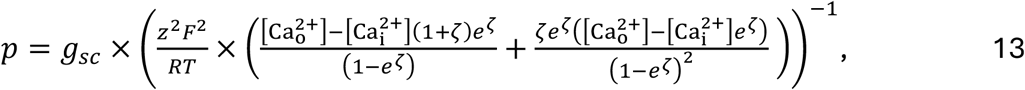

where 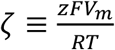 and *V* = -70 *mV*.

## Channel gating

The voltage-dependent equilibrium for each gate (x = m or h) and its time-dependent and steady state open probability are expressed below. All gates must open for the channel to conduct a current.

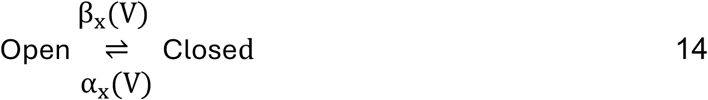

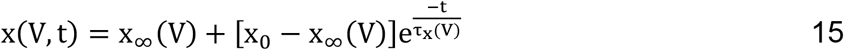

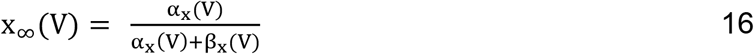

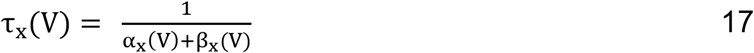

Where *α* and *β* are voltage-dependent transition rates and *τ* denotes the time constant of activation or inactivation.

## Thermodynamic model of rate constants

We employ the thermodynamic model of voltage dependent rate constants described by Destexhe and Huguenard.^*51*^ Briefly, each transition rate, r = *α*_*x*_, *β*_*x*_ is described by a Boltzmann distribution, where *ΔG* is the free energy barrier associated with the transition, *R* is the gas constant, and *T* is the temperature.

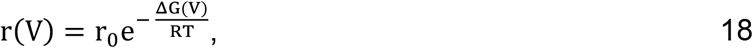

where Δ*G* is expressed as a nonlinear function of voltage.

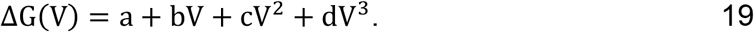

Defining *V*_*x*_ as the voltage at which *α*_*x*_ *= β*_*x*_, we arrive at the following rate expressions for the *m* and *h* gates (see *Supplementary Note 2* for a definition of *m* and *h*).

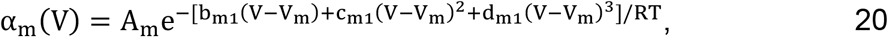

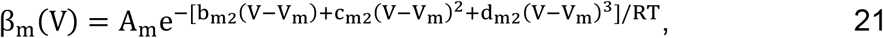

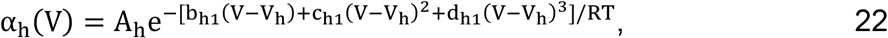

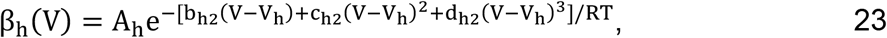

Using the parameters reported in Destexhe *et al*. as a starting point, we further refined the parameter values to fit our macroscopic Ca_V_3.2 currents (see *Supplementary Table 2* and Fig. S6).

**Supplementary Note 3. Stochastic simulation of single-channel trajectories**. We employed the Gillespie algorithm^*52*^ to simulate single-channel trajectories using thermodynamic models of transition rates fit to our ensemble-average currents. Our implementation of the algorithm is as follows:

Define voltage-dependent transition rates (Eq. 14), α_x_(V), β,_x_(V). Assume equilibrium at voltage V = V_hold_ prior to the first time-step, t_1_.

1. assign an initial open probability P_O_(t_0_)

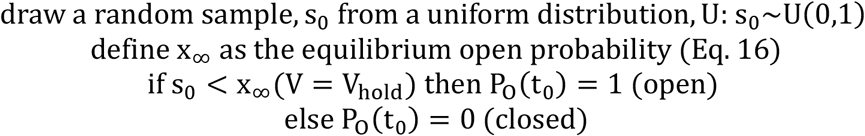
2. instantaneously step to V = V_test_
3. sample the gate dwell time, τ in the t_0_ state (open or closed) for P_O_(t_0_) = 1 (open)

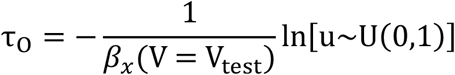

for P_O_(t_0_) = 0 (closed)

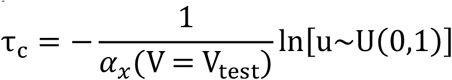
4. sample the following state for P_O_(t_0_)=1

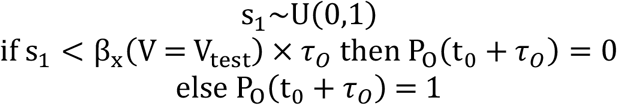

for P_O_(t_0_) = 0

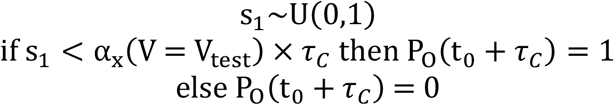

repeat for t ≤ 45 ms

We estimated the distribution of charge influx from simulated single-channel trajectories by integrating Eq. 12 over *t*_V_ (Fig. 5D, S6).

**Supplementary Table 1.**
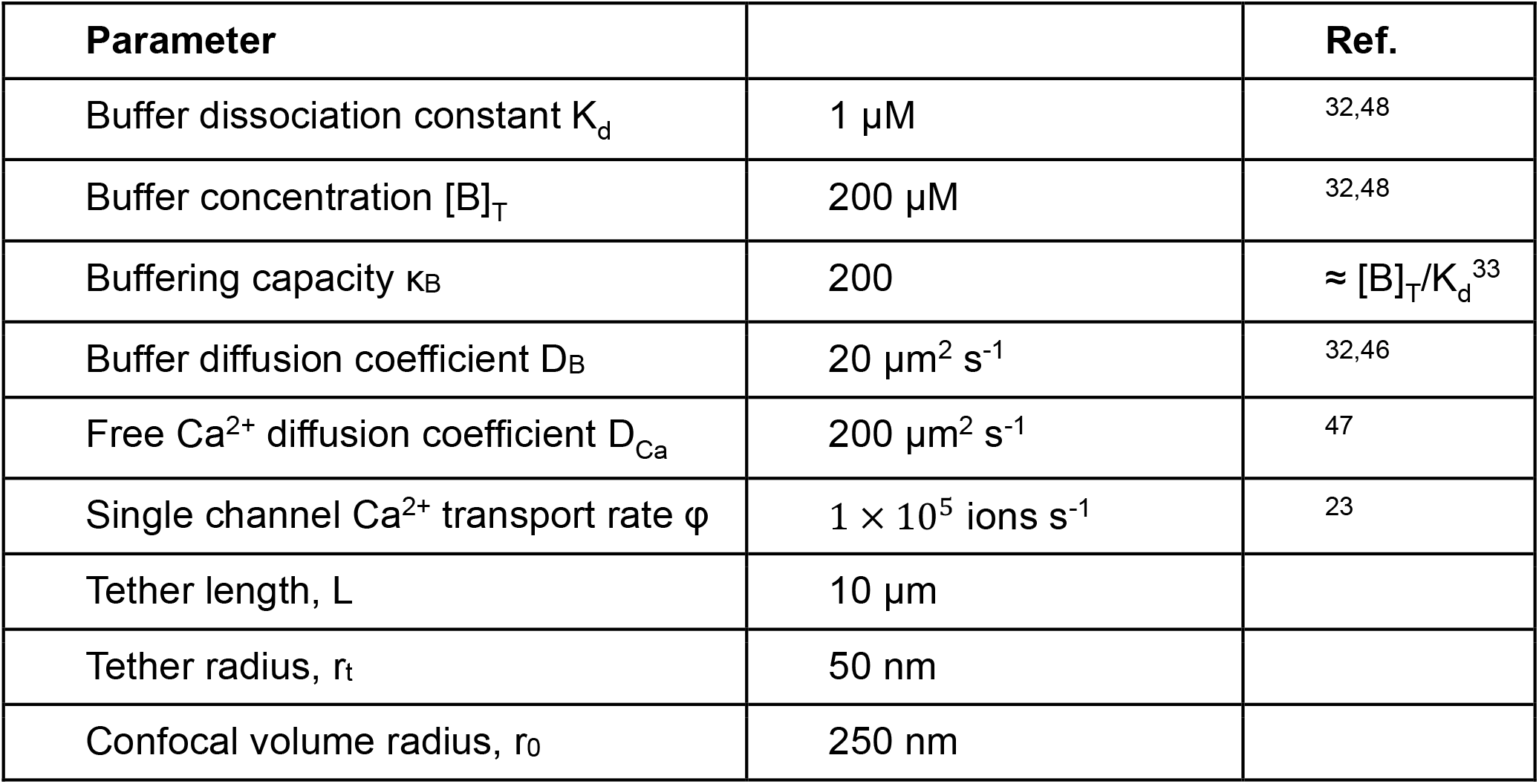
Parameters for modeling single-channel Ca^2+^ domains.

| Parameter |  | Ref. |
| --- | --- | --- |
| Buffer dissociation constant $K_d$ | 1 $\mu\text{M}$ | 32,48 |
| Buffer concentration $[B]_T$ | 200 $\mu\text{M}$ | 32,48 |
| Buffering capacity $\kappa_B$ | 200 | $\approx [B]_T/K_d^{33}$ |
| Buffer diffusion coefficient $D_B$ | 20 $\mu\text{m}^2 \text{s}^{-1}$ | 32,46 |
| Free Ca<sup>2+</sup> diffusion coefficient $D_{Ca}$ | 200 $\mu\text{m}^2 \text{s}^{-1}$ | 47 |
| Single channel Ca<sup>2+</sup> transport rate $\varphi$ | $1 \times 10^5 \text{ ions s}^{-1}$ | 23 |
| Tether length, $L$ | 10 $\mu\text{m}$ | |
| Tether radius, $r_t$ | 50 nm | |
| Confocal volume radius, $r_0$ | 250 nm | |

**Supplementary Table 2.** Fit of model parameters to macroscopic currents. See *Supplementary Note 2* for a description of the gating model. T = 32 °C

| m-gate parameters |  | h-gate parameters |  |
| --- | --- | --- | --- |
| $V_m$ | -49 mV | $V_h$ | -69 mV |
| $A_m$ | 0.093 ms<sup>-1</sup> | $A_h$ | 2.7e-3 ms<sup>-1</sup> |
| $b_{m1}$ | -340 J mV<sup>-1</sup> | $b_{h1}$ | 150 J mV<sup>-1</sup> |
| $c_{m1}$ | 3.6 J mV<sup>-2</sup> | $c_{h1}$ | 6.5 J mV<sup>-2</sup> |
| $d_{m1}$ | 5.4e-4 J mV<sup>-3</sup> | $d_{h1}$ | 0.062 J mV<sup>-3</sup> |
| $b_{m2}$ | 130 J mV<sup>-1</sup> | $b_{h2}$ | -470 J mV<sup>-1</sup> |
| $c_{m2}$ | 3.8 J mV<sup>-2</sup> | $c_{h2}$ | 7.0 J mV<sup>-2</sup> |
| $d_{m2}$ | 0.018 J mV<sup>-3</sup> | $d_{h2}$ | -0.057 J mV<sup>-3</sup> |

**Supplementary Figure 1.**
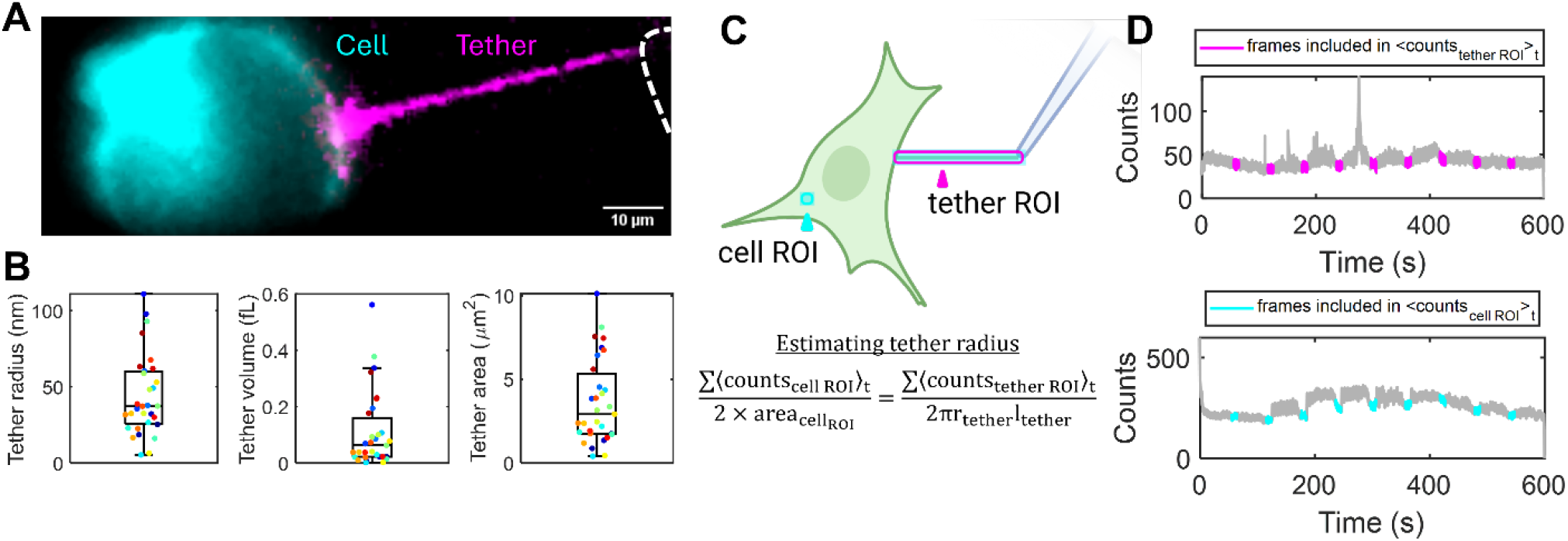
Characterization of tether geometry. **A**. Fluorescence image of a membrane tether extracted from a HEK293 cell expressing lck-jGCaMP8f and Ca_V_3.2. Image is a composite of two pictures, one with tether-only illumination (purple) and one with cell body illumination (cyan). Tether-only contrast is enhanced to aid visualization. **B**. Distribution of tether radii (left, n = 31 tethers, median 37.2 nm), volumes (middle, median 62.7 aL), and surface areas (right, median 2.91 μm^2^) estimated using baseline fluorescence counts/membrane area from cell body lck-jGCaMP8f or GCaMP6s-CAAX and assuming cylindrical tether geometry (box: median, 25^th^ and 75^th^ percentiles, extrema). **C**. Procedure for estimating radii of tethers. Excitation light (light blue) is restricted to the tether (magenta) and a small patch of the cell membrane (cyan). **D**. Cell and tether GECI fluorescence, recorded in the absence of depolarizing stimulus (cyan and magenta timepoints), are averaged in time and summed over region of interest (ROI). Bottom: cell ROI counts per ROI area (accounting for excitation of top and bottom membrane patches) provides a count density, which is used to estimate tether radius.

**Supplementary Figure 2.**
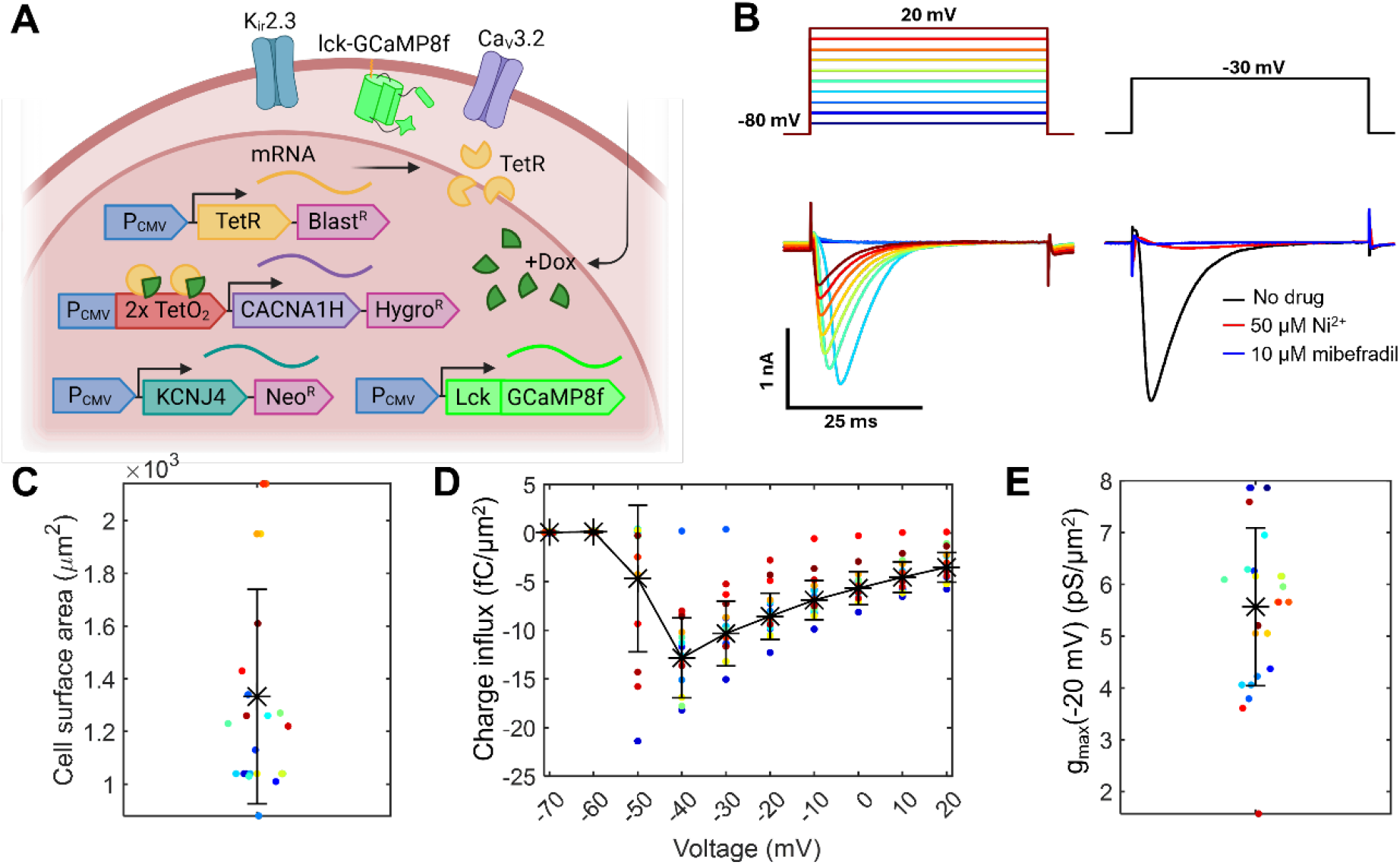
Electrophysiology of HEK cells expressing Ca_V_3.2. **A**. HEK293 cell line stably incorporated genes encoding Tet repressor (TetR), tetracycline-inducible Ca_V_3.2 pore-forming subunit *CACNA1H*, and constitutive K_ir_2.3 inward-rectifier potassium channel *KCNJ4*. Membrane-targeted GECI (lck-jGCaMP8f shown here) was expressed by transient transfection. **B**. Representative inward currents (bottom, left) in response to depolarizing voltage pulses (top). Ca_V_3.2 channel blockers suppressed these currents (bottom, right). **C**. Distribution of cell surface areas, calculated from whole cell capacitance, assuming a specific membrane capacitance of 1 μF/cm^2^ (1332 ± 406 μm^2^, mean ± s.d., n = 17 cells). **D**. Distribution of charge influx densities. Cells were treated with 1.5 ng/mL doxycycline and measured at 24-48 hr post-induction. Charge influx density was calculated via the time integral of whole-cell inward currents and dividing by the cell surface area (mean ± s.d., n = 17 cells). **E**. Distribution of conductance densities for the cells analyzed in C-D. Calculated by taking the maximal inward current measured at -20 mV and dividing by the Ca^2+^ driving force (5.57 ± 1.52 pS/μm^2^, mean ± s.d., n = 17 cells). Multiplying the average conductance density by the median tether area (2.9 μm^2^, Fig. S1), and dividing by prior estimates of channel conductance^*23*^ (g = 1.7 pS) and predicted maximum channel open probability (P_o_ = 0.82) yields an estimate of ∼12 channels per tether.

**Supplementary Figure 3.**
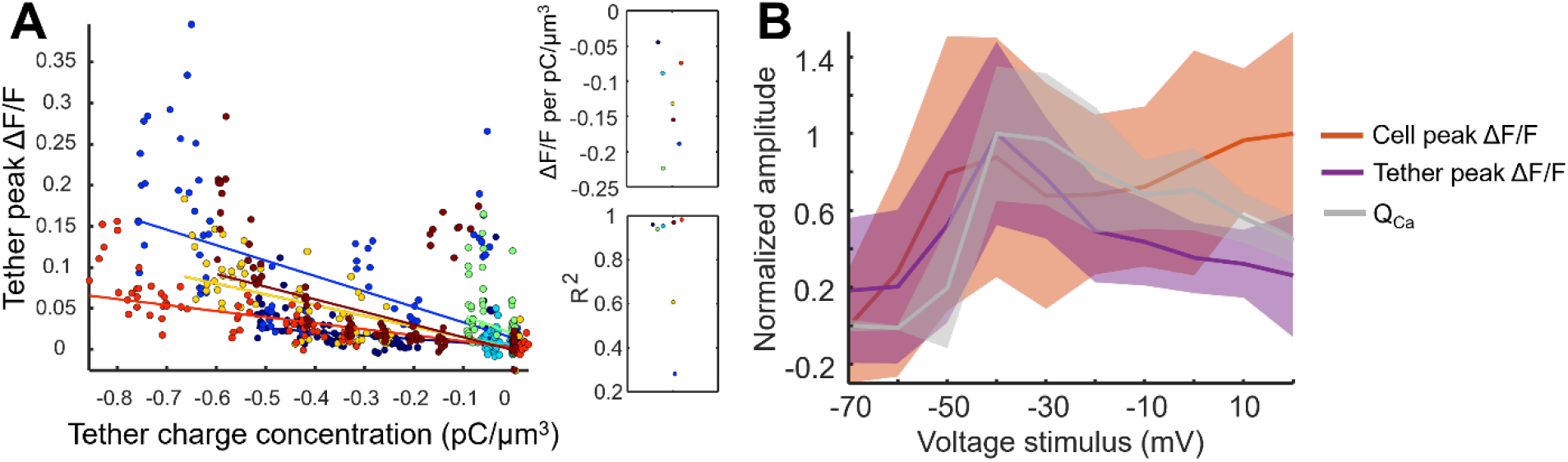
Tether Ca^2+^, reported by GCaMP6s-CAAX, correlates with ensemble charge transport. **A**. Peak (ΔF/F)^t^ amplitude of tether GCaMP6s-CAAX fluorescence transients as a function of predicted tether charge concentration, 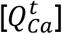, and linear fits. Inset: slope (top) and R^2^ (bottom) from linear fits of n = 7 tethers. **B**. Voltage dependence of stimulus-evoked charge transport (gray), tether (purple) and cell (orange) peak ΔF/F amplitudes averaged across n = 7 GCaMP6s-CAAX cell-tether pairs. Traces are normalized for each tether-cell pair before averaging and shaded to show std. dev.

**Supplementary Figure 4.**
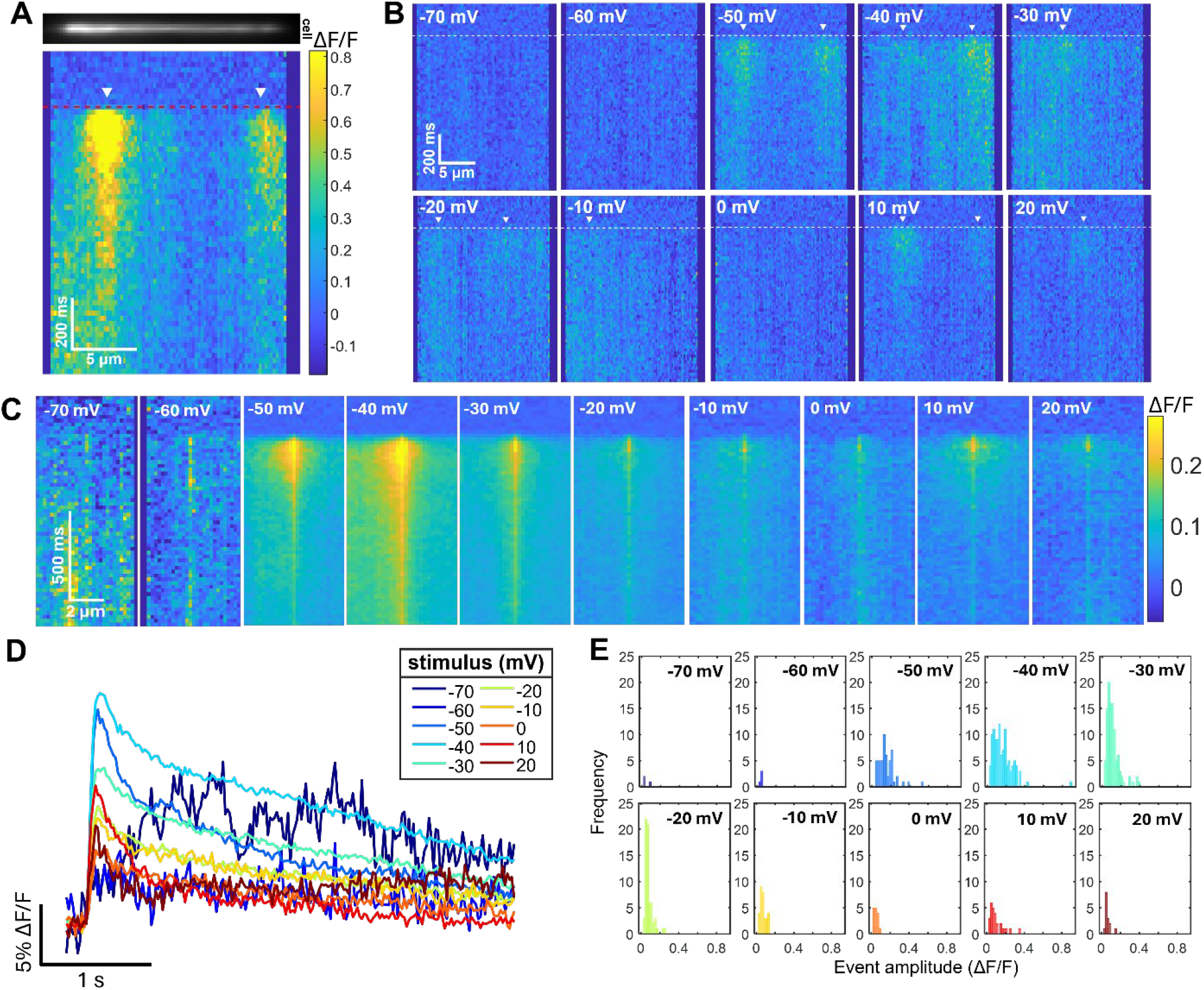
Spatiotemporal structure and statistics of Ca^2+^ events reported by GCaMP6s-CAAX indicator. **A**. Epifluorescence image of a tether above a kymograph for a 45 ms voltage step from -80 mV to -30 mV. Red dashed line indicates step onset. White arrowheads indicate discrete Ca^2+^ events. **B**. Kymographs of tether Ca^2+^-dependent fluorescence in response to voltage steps from -80 mV to between -70 mV and +20 mV. White dashed lie indicates step onset. Data shown for single-trial responses of one tether. **C**. Stimulus-triggered average kymographs of spatially isolated tether Ca^2+^ events (n = 3-106 events per voltage, 7 tethers). **D**. Timecourse of events in (C), averaged over ±2.2 μm from the fluorescence peak. **E**. Histogram of Ca^2+^ event amplitudes at each stimulus voltage (n = 3-106 events per voltage, 7 tethers). Event amplitudes are taken from spatiotemporally filtered kymographs.

**Supplementary Figure 5.**
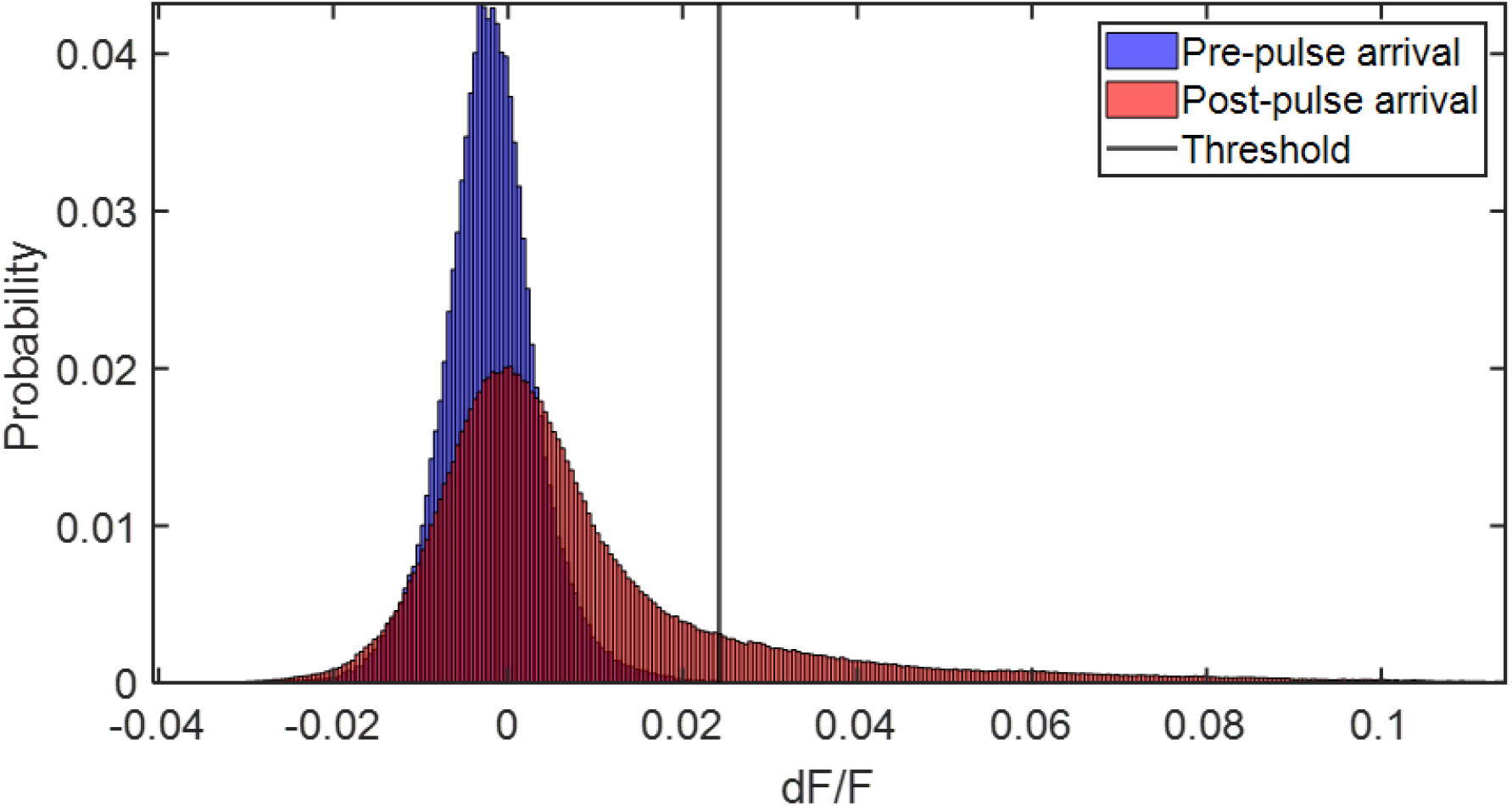
Ca^2+^ event detection threshold in tethers from HEK cells expressing lck-jGCaMP8f. Distribution of pixel intensities within pre-stimulus window (V_m_ = -80 mV, blue) and all pixels within post-stimulus window (red, V_m_ = -70 to 20 mV, includes Ca^2+^ event and non-event pixels) for one lck-jGCaMP8f expressing tether. Black line indicates threshold used to segment Ca^2+^ events. Threshold calculation described in *Methods*.

**Supplementary Figure 6.**
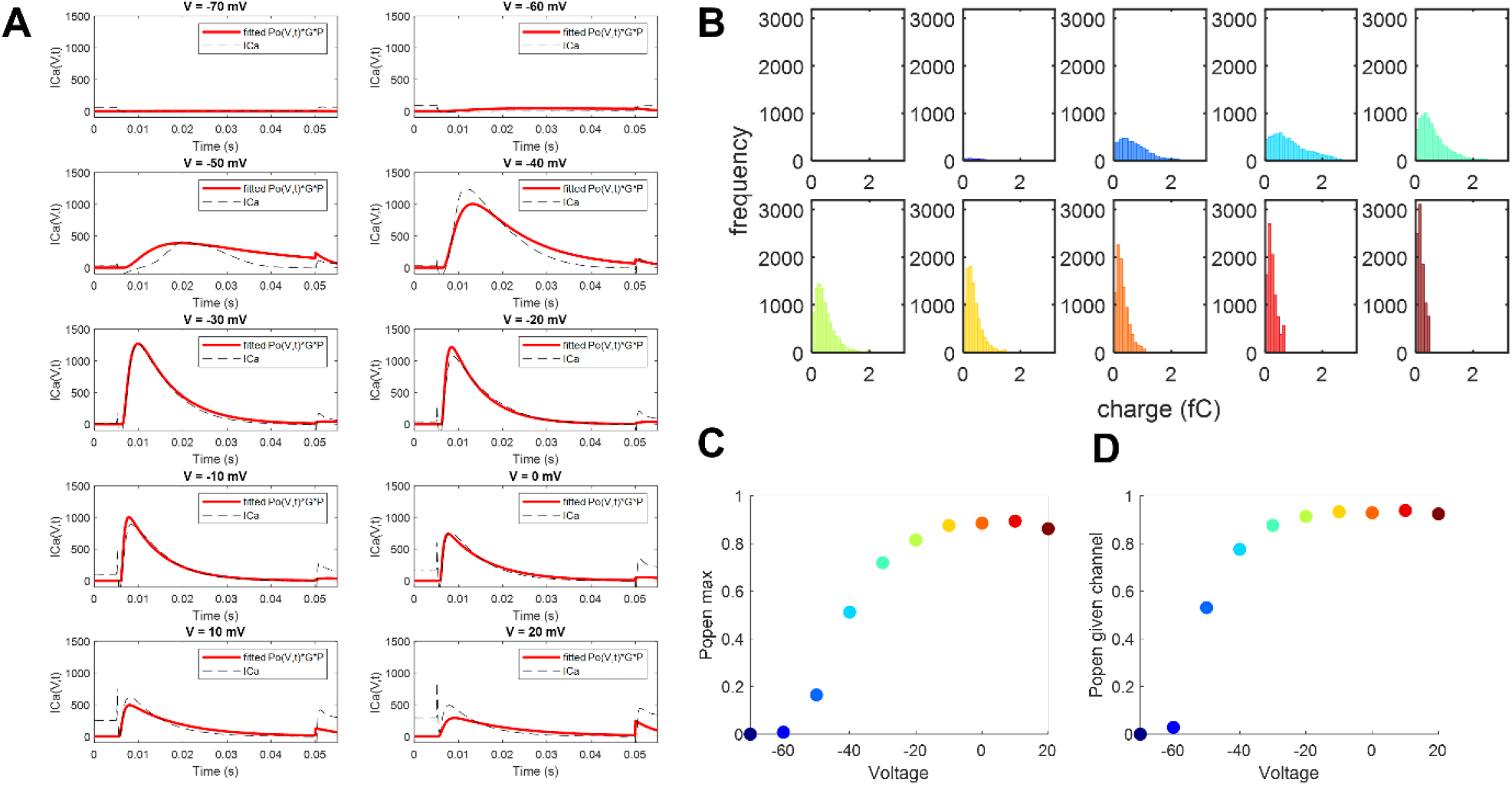
Refinement of gating model and simulation of Ca_V_3.2 channel trajectories. **A**. Comparison of measured macroscopic currents (dashed black line, average of n = 17 cells) and *I*_*Ca*_ (Eq. 11) simulated using a nonlinear thermodynamic model of transition rates (Eq. 20-23) with fitted parameters (red trace, *Supplementary Table 2*). **B**. Histogram of total charge conducted during simulated 45 ms single-channel trajectories (see *Supplementary Note 3* for a description of the stochastic simulation). **C**. Peak open probability during 45 ms stimulation for an ensemble of simulated channels. **D**. Predicted probability of a given channel opening at least once during 45 ms stimulation. N = 10^4^ channel trajectories simulated for B-D.

**Supplementary Figure 7.**
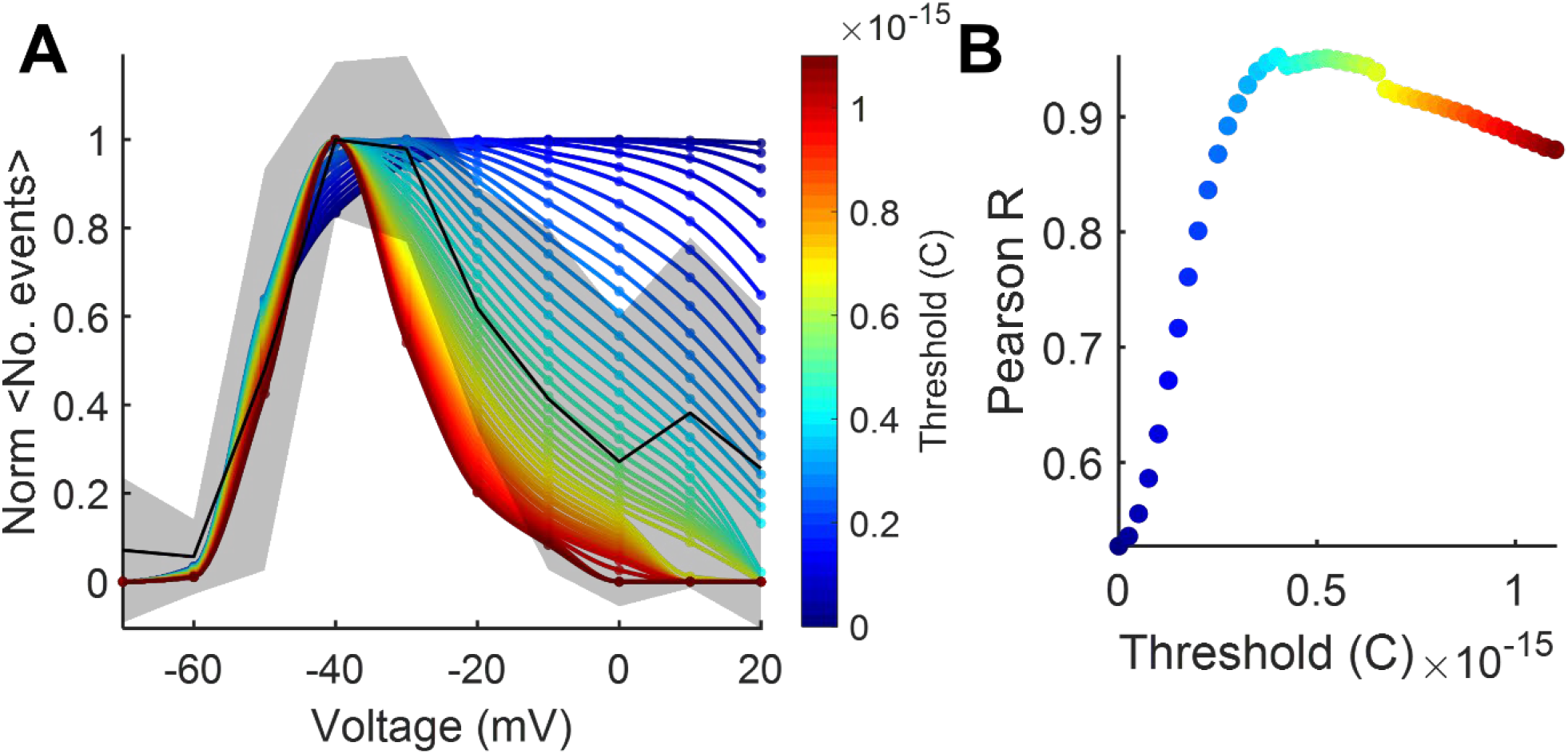
Detection threshold for Ca^2+^ events reported by GCaMP6s-CAAX fluorescence. **A**. Mean number of events vs. voltage (normalized, black, shading std. dev. of n = 7 tethers) and stochastic simulations of the probability of observing a single-channel event (colors). Color bar indicates simulated charge detection threshold. **B**. Pearson correlation between the frequency of observed events (A, black) and the predicted single-channel event probability (A, colors) as function of detection threshold (maximum R-value at 0.4 fC threshold).

